# Staining-Free, In-Flow Enumeration of Tumor Cells in Blood Using Digital Holographic Microscopy and Deep Learning

**DOI:** 10.1101/2022.05.01.490222

**Authors:** Anirudh Gangadhar, Hamed Sari-Sarraf, Siva A. Vanapalli

**Affiliations:** Department of Chemical Engineering, Texas Tech University, Lubbock, TX 79409; Department of Electrical and Computer Engineering, Texas Tech University, Lubbock, TX 79409

## Abstract

Currently, detection of circulating tumor cells (CTCs) in cancer patient blood samples relies on immunostaining, which does not provide access to live CTCs, limiting the breadth of CTC-based applications. As a first step to address this limitation, here, we demonstrate staining-free enumeration of tumor cells spiked into lysed blood samples using digital holographic microscopy (DHM), microfluidics and machine learning (ML). A 3D-printed module for laser assembly was developed to simplify the optical set up for holographic imaging of cells flowing through a sheath-based microfluidic device. Computational reconstruction of the holograms was performed to localize the cells in 3D and obtain the plane of best focus images to train deep learning models. First, we evaluated the classification performance of two convolutional neural networks (CNNs): ResNet-50 and a custom-designed shallow Network dubbed s-Net. The accuracy, sensitivity and specificity of these networks were found to range from 97.08% and 99.32%. Upon selecting the s-Net due to its simple architecture and low computational burden, we formulated a decision gating strategy to significantly lower the false positive rate (FPR). By applying an optimized decision threshold to mixed samples prepared *in silico*, the FPR was reduced from 1×10^−2^ to 2.77×10^−4^. Finally, the developed DHM-ML framework was successfully applied to enumerate spiked MCF-7 breast cancer cells from lysed blood samples containing a background of white blood cells (WBCs). We conclude by discussing the advances that need to be made to translate the DHM-ML approach to staining-free enumeration of CTCs in cancer patient blood samples.

## 1. Introduction

Circulating tumor cells (CTCs) shed from primary tumors have been identified in the blood of cancer patients^1^. Given their critical role in the metastatic cascade, these rare cells serve as novel clinical biomarkers to assess the tumor burden enabling effective monitoring of therapeutic treatment strategies^2–4^. This has led to numerous efforts to develop methods to identify and characterize these information-laden cells. In comparison to traditional tissue biopsies, these blood-based “*liquid biopsy*” approaches are minimally invasive and can be carried out routinely enabling more frequent monitoring of CTCs^5^.

CTCs occur at extremely low frequencies in the blood (~1-100 CTCs/10^9^ blood cells)^6–8^, motivating the development of methods to isolate them. Two main types of separation strategies are employed to isolate CTCs. In the affinity-based approach, specific antibodies are immobilized onto solid substrates, whereupon CTCs expressing the target antigen are selectively captured and recovered for subsequent downstream processing^9–12^. In contrast, label-free methods take advantage of the differences in biophysical properties such as size^13–15^, deformability^16^, dielectric properties^17,18^ and density^19^ between CTCs and blood cells. Both class of CTC isolation methods do not offer 100% purity, resulting in a mixture of CTCs and nucleated white blood cells (WBCs), with the ratios of CTCs to WBCs ranging from 1:100 – 1:1000 per mL after enrichment^20–22^.

The presence of background WBCs in the isolated sample makes it necessary to perform immunostaining – the current gold standard to identify CTCs using antibodies targeting CTC-specific antigens. However, the use of immunostaining limits the full translational utility of CTCs. For example, there is growing interest in ex-vivo culture of CTCs for biomarker and drug discovery^23–25^ and the bottleneck here is to know which patient samples have sufficient CTCs for culture expansion since blood samples may not even contain CTCs or have very low counts depending on cancer type, stage and treatment course^26,27^. Given that the immunostaining process kills CTCs, it becomes impossible to know apriori which patient samples are suitable for CTC culture. Therefore, it is useful to develop a staining-free quick screening tool that will enable rapid assessment of CTC load allowing informed decisions on the type of downstream assays that could be performed with live CTCs isolated from each cancer patient blood sample.

Staining-free approaches to detect CTCs could be potentially developed from bright-field images of CTCs and WBCs^28^. However, marker-free CTC isolation methods (e.g. Vortex HT chip^29^ and Labyrinth chip^30^) often generate the rare cells in large sample volumes, requiring spinning down of the suspension to a localized spot on glass side to facilitate imaging, but this additional spin step can result in significant cell losses^31–33^. Addressing this issue, previously, a study from our laboratory has shown that application of in-line digital holographic microscopy (DHM) to cells flowing in a microchannel, can process large sample volumes and fingerprint individual cells^34^, without the need to concentrate the sample. This flow-through DHM approach is attractive since 3D information of flowing objects is encoded onto a 2D image recorded on the camera sensor. The resulting interference patterns or holograms can then be computationally reconstructed to obtain 3D positional information, size and intensity features that can be used to differentiate cell types^35,36^.

Unlike our previous work where the in-line DHM optical set up was complex and engineered features from hologram processing were used to differentiate cell types, here (i) we simplified the optical set up to develop a portable DHM (pDHM) module that is compact and can be instrumented to a standard inverted microscope (ii) we implemented convolutional neural network (CNN) on the in-focus images extracted from the holograms to train a machine learning (ML) model that can distinguish and enumerate cancer cells and WBCs in flow. This DHM-ML-assisted approach to enumerate cancer cells among a background of WBCs, could eliminate the complex and time-consuming multi-step immunostaining procedure, enabling rational decision-making on the types of downstream assays that can be pursued with blood samples of cancer patients.

Our paper is organized as follows: We begin with a description of our pDHM approach in Sec. 2.1, followed by detailing the analysis workflow developed for flow-based interrogation of tumor cells in Sec. 2.2. Next, in Sec. 2.3, we compare the performance of two ML networks: ResNet-50 and a custom-built shallow network (s-Net) in classifying in vitro MCF-7 breast cancer cells from WBCs using reconstructed plane of best focus (PoBF) holographic images. In Sec. 2.4, the effectiveness of the decision threshold is further tested on artificial mixed datasets prepared from the pure population images of both cell types. In Sec. 2.5, we apply our pDHM-ML framework to enumerate breast cancer cells spiked into lysed blood at various target concentrations. In Sec. 2.6, we contrast our work against prior studies. Finally, in Sec. 3, we discuss the implications of our findings and the challenges that need to be addressed to translate our work to actual CTCs in cancer patient blood.

## 2. Results

### 2.1 Portable digital holographic microscopy (pDHM) module

Digital holographic microscopy (DHM) has been widely used as a quantitative imaging tool in various cell-based applications such as morphology characterization using optical properties like intrinsic refractive index^37,38^, mapping 3D trajectories in flow^39,40^ and label-free detection of diseases such as cancer^41–43^. Two types of DHM configurations have been commonly utilized, in-line and off-axis, so named depending on whether the light source, sample and recording medium share the same optical axis or not^39,44–46^. A common drawback is that these optical set ups can be complex, requiring precision alignment of multiple components, making it challenging to easily translate this powerful technique to laboratories.

Addressing this need, we have developed the pDHM module that sits on the stage of a standard inverted microscope (Fig. 1a). It consists of two components: (1) a laser torch which contains within it a laser diode, a collimating lens and an automatic power control circuit and (2) a 3D-printed assembly to mount and position the torch. The emitted laser beam has a wavelength of 635 nm and a minimum diameter of 3 mm. The source-to-sample distance is about 45 mm while the maximum distance between the sample and the focal plane of the microscope objective is 0.53 mm.

**Figure 1.**
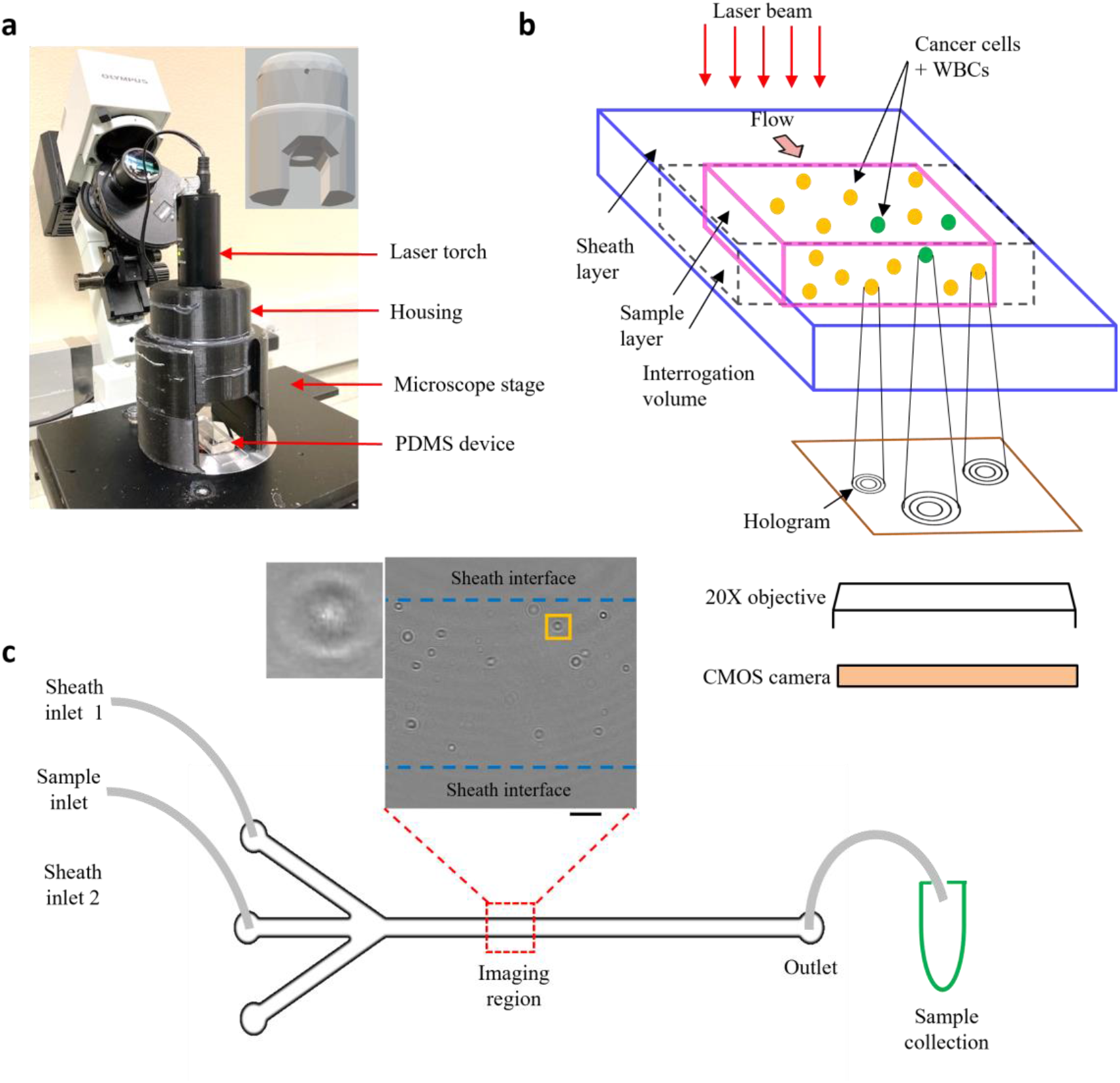
Experimental setup. (a) Portable holographic assembly. The laser torch is mounted on a movable housing. Inset shows the 3D design of the housing, the circular aperture allows the laser beam to pass through, illuminating the sample coherently; (b) Schematic showing in-flow holograms of a mixed sample containing MCF7 cells and WBCs imaged with a 20X microscope objective and recorded digitally, using a CMOS, high-speed camera; (c) PDMS-based sheathing device used for flowing the sample contains two inlet channels for entry of the sheath fluid (PBS) and a central inlet for sample injection. Imaging is performed in a region near the center of the channel far away from the entrance. Inset shows a representative cleaned hologram recorded using the pDHM setup, scalebar is 100 *μm*. By imposing a sheath flow, the cells are confined to a region away from the channel walls. Post-DHM imaging, sample is collected from the device outlet for subsequent fluorescence imaging to obtain the ground truth counts.

The schematic in Fig. 1b illustrates the hologram recording process. Cells flowing through a microfluidic channel scatter the incident laser light. The scattered (object) wave interacts with the unobstructed (reference) wave to form interference fringes or holograms at the focal plane of the microscope objective. These are subsequently recorded digitally on a CMOS sensor using a high-speed camera. Since the source, sample and detector are located along the same optical axis, holographic imaging is performed using the in-line configuration. In the in-line mode, reconstructed intensity fields are obtained from the corresponding amplitudes while the phase information gets corrupted due to the twin-image artefact, requiring additional recovery strategies^47–49^.

Unlike, our previous study, here we modified the microchannel geometry to include a trifurcated inlet (Fig. 1c). The microchannel had a width (W) and height (H) of W × H = 800 *μm* × 330 *μm* respectively. The trifurcated inlet was used to introduce a sheathing flow to overcome two main issues: (1) undesirable interference fringes from channel sidewalls and (2) cells close to the wall move very slowly, creating multiple instances in the hologram recordings. These multiple cell counts could introduce large errors especially for the near-wall cells. By controlling the flow rate of the sheath fluid relative to the sample flow rate, the cell suspension was confined to central width of ≈ 500 *μm*, and a ≈ 150 *μm* of sheath fluid on either side. At an optical resolution of 1 *μm* per pixel, a single hologram corresponds to an image volume of 500 *μm* × 800 *μm* × 330 *μm* ≈ 0.13 *μL* of sample. To remove multiple cell counts, a frame rate of 420 fps was chosen such that the fastest moving cell only appeared in a single frame. Typically, it was observed that cells appeared in a maximum of 2-3 sequential frames. These cells were mostly located near the sample-sheath interface wherein, the parabolic fluid velocity profile caused them to move much slower as compared to cells near the channel centerline. At the selected frame rate, 10100 holograms were captured so that tumor cells could be enumerated from 1 mL of sample volume flowing through the microchannel.

### 2.2 Basic workflow of DHM-ML detection of cancer cells

To implement the DHM-ML strategy, we used MCF-7 breast cancer cells as a surrogate for CTCs recognizing that some characteristics of patient-derived CTCs could be different from that of MCF-7 cells. Blood was processed using lysis buffer to eliminate red blood cells and centrifuged to isolate WBCs. Holograms were obtained of pure populations of MCF-7s as well as WBCs that were used for selection of ML models. Mixed samples containing MCF-7s spiked into WBCs were used to test the performance of the optimal ML model. The MCF-7 cells were labeled with a fluorescent marker to establish ground-truth validation of counts obtained from the DHM-ML strategy.

The computational analysis workflow for the DHM-ML strategy is shown in Fig. 2a. To profile every cell in the field of view (FOV), the acquired raw holograms are first denoised by subtracting each raw hologram with an object-free background, estimated as the average of 10,100 raw holograms. Subsequently, numerical reconstruction is performed on the resultant hologram to localize in 3D all the cells in the image volume. From the reconstructed image stack, the 2D center of each cell is identified initially which enables its detection in the denoised hologram (Fig. 2b). Equipped with this information, axial intensity maps are generated for each cell which allows us to obtain the plane of best focus (PoBF) or in-focus image (Fig. 2c). Examples of PoBF images for MCF-7s and WBCs are shown in Fig. 2d. Our methodology for background subtraction, reconstruction of holograms and PoBF selection has been described in detail previously^34,45^.

**Figure 2.**
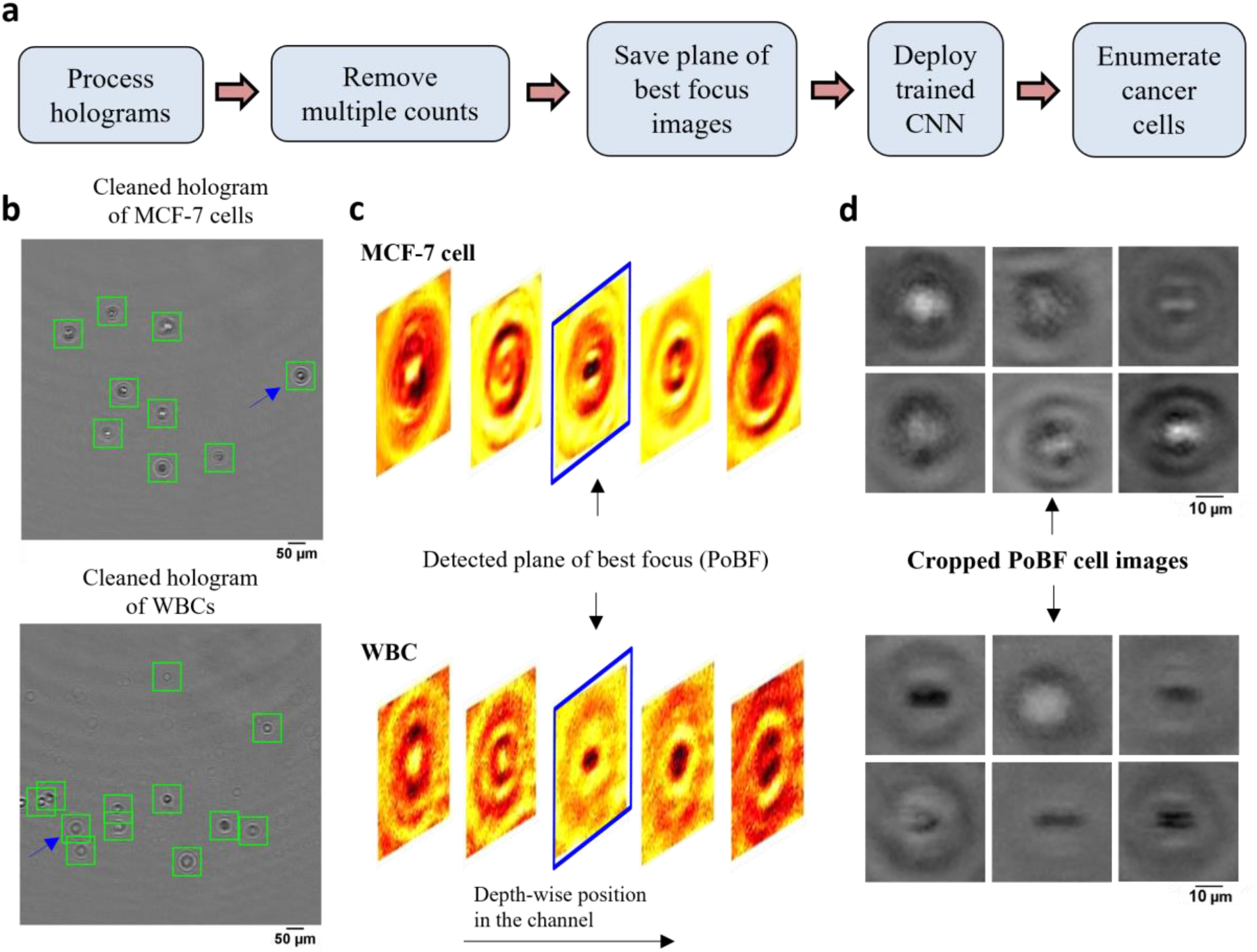
DHM-ML workflow including extraction of plane of best focus images of cells. (a) DHM-ML analysis workflow; (b) Cells detected and localized as shown by the green squares are overlaid on the cleaned holograms of pure cell populations; (c) Full volume numerical reconstruction allows 3D localization of cells, enabling detection of the plane of best focus (PoBF). Reconstructed images of the cell highlighted by the blue arrow in (a) are shown at various axial distances. The image highlighted in blue shows the detected plane of best focus (PoBF); (d) Examples of cropped, 36×36 PoBF images are shown for pure MCF-7 cells (top) as well as WBCs (bottom).

Next, multiple counts of cells are removed since the flow is laminar where the motion of cells is predominantly in the streamwise direction and the cross-stream (*y*) and axial (*z*) velocity components can be neglected. This allows us to use their detected *y, z* positions to remove multiple counts. Cells were identified and eliminated if the following conditions were met: |*y*_*q,i*+1_ − *y_p,i_* | ≤ 3 *μm* and |*z*_*q,i*+1_ − *z_p,i_*| ≤ 50 *μm*. These criteria were verified by manually tracking the detected *y, z* positions of about 100 MCF-7 cells as well as WBCs selected at random. Interestingly, we found appreciable changes in the axial positions of cancer cells between two successive frames, consequently, a relatively lenient *z* criterion is applied.

After removing the multiple counts, the PoBF images (Fig. 2d) are cropped to 36 × 36 pixel sub-images using the centroid location of the cells. These sub-images are fed to the trained deep learning model which classifies them as MCF-7 or WBC. Applying an optimized decision gating on the predictions, we obtain enumerated counts of MCF-7 cells in the analyzed sample. It is worth noting that the entire computational analysis pipeline is automated and does not require any supervision.

### 2.3 ML Model Selection: ResNet versus s-Net

In this section, we seek to evaluate the performance of DL models in classifying MCF-7 breast cancer cells from WBCs using holographic reconstructed in-focus images. To achieve this, we tested two CNN architectures: (1) ResNet-50, a popular choice for image-based classification tasks^28^ and (2) a custom-built, shallow network, s-Net, with significantly fewer parameters. Advantages of using the latter mainly include shorter training and testing times without the need for high computational power. We evaluated the classification performance of the s-Net and compared it with that of ResNet-50.

The s-Net architecture is shown schematically in Fig. 3a. A 36×36 PoBF cell image is fed to the network as input. Three convolutional layers make up the backbone with a filter size of 3×3 and a stride of 1. The number of filters increases from 8 to 32 and “same” padding is used to ensure that image size is conserved before and after the convolution. This is followed by batch normalization and ReLU activation. The output image is passed to a max pooling layer where it is down sampled using a window size and stride of 2. The third convolutional layer is succeeded by a flattened fully connected (FC) layer containing 2592 nodes. Finally, a Softmax layer produces the class (MCF-7, WBC) probabilities which result in a prediction. The activations and number of learnable parameters (weights + biases) are tabulated in Fig. 3b.

**Figure 3.**
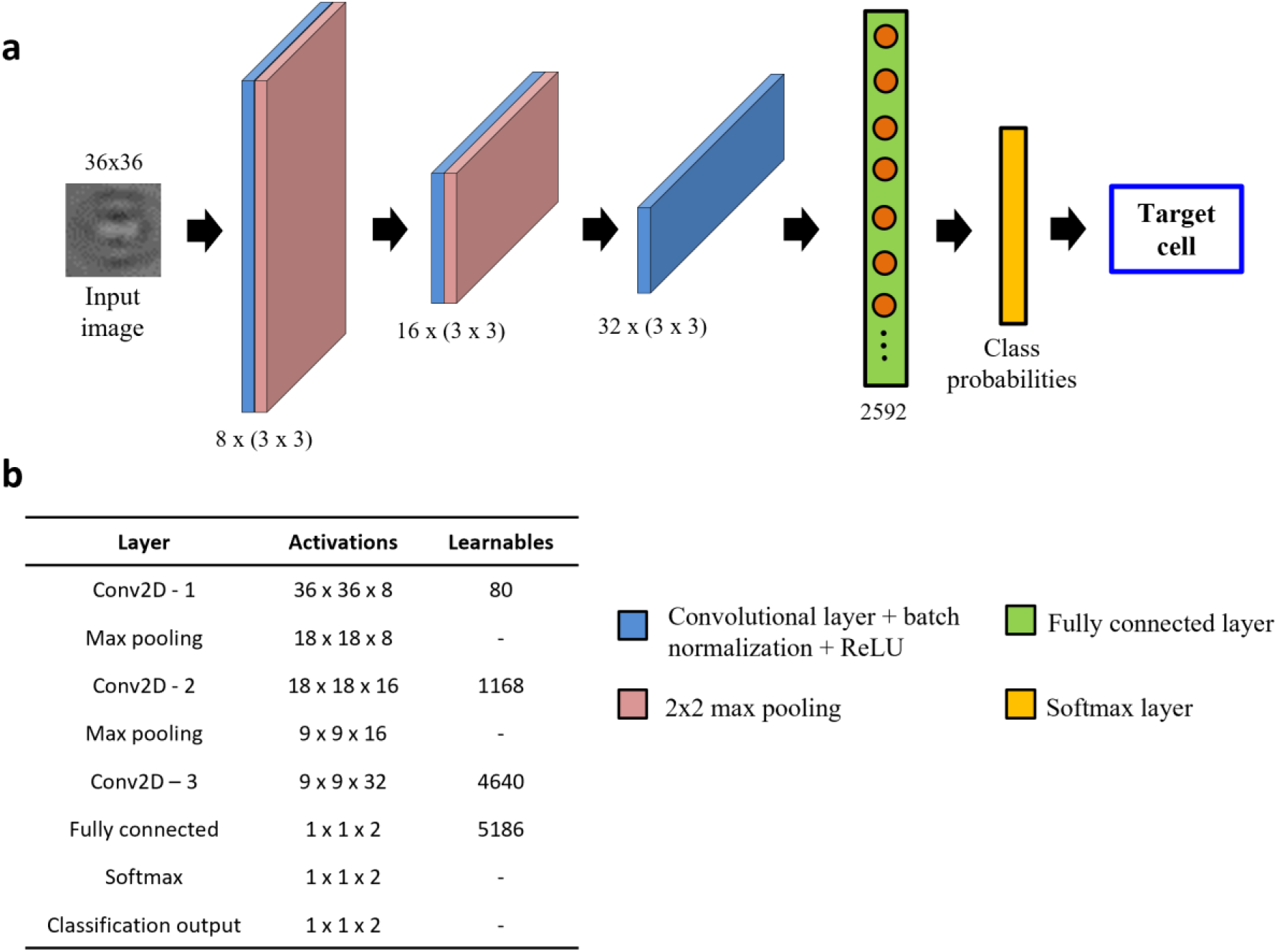
Architecture of custom-built s-Net. (a) Network architecture consisting of 8 layers: 3 convolutional, 2 max pooling, 1 fully connected, Softmax and output classification; (b) Table showing the number of activations and learnables (sum of weights and biases) in each layer. The Network learns parameters only in the convolutional and fully connected (FC) layers.

For model evaluation, training and testing set images were generated from pure population experiments, i.e., only MCF-7 or WBC images. A total of 52,340 images were used per class. A 70/30 split was used to randomly generate the training and testing datasets which yielded 36,638 and 15,702 class images respectively. We used a minibatch size of 32 and the number of epochs was set at 20. To compute the network weights, Adam optimizer with an initial learning rate of 10^−3^ was selected. To ensure consistency, the same training parameters were used for both models.

Before discussing the CNN performance, it is important to clearly define these measures along with some ML-specific terminologies: True positives (TPs) refer to MCF-7 cells classified correctly; True negatives (TNs) refer to WBCs classified correctly; False positives (FPs) refer to WBCs incorrectly classified as MCF-7 cells; False negatives (FNs) refer to MCF-7 cells incorrectly classified as WBCs. In terms of performance measures, we define Accuracy A = (TP+TN)/(TP+TN+FP+FN); Sensitivity Se = TP/(TP+FN) also known as true positive rate (TPR); Specificity Sp = TN/(TN+FP) also known as true negative rate (TNR); and False positive rate (FPR): FPR = FP/(FP+TN). Fig. 4a shows the confusion matrices computed on the testing dataset for both networks. Performance of the trained networks was evaluated by computing the accuracy, sensitivity and specificity on the testing dataset (Fig. 4b). Overall, we find that both networks perform reasonably well.

**Figure 4.**
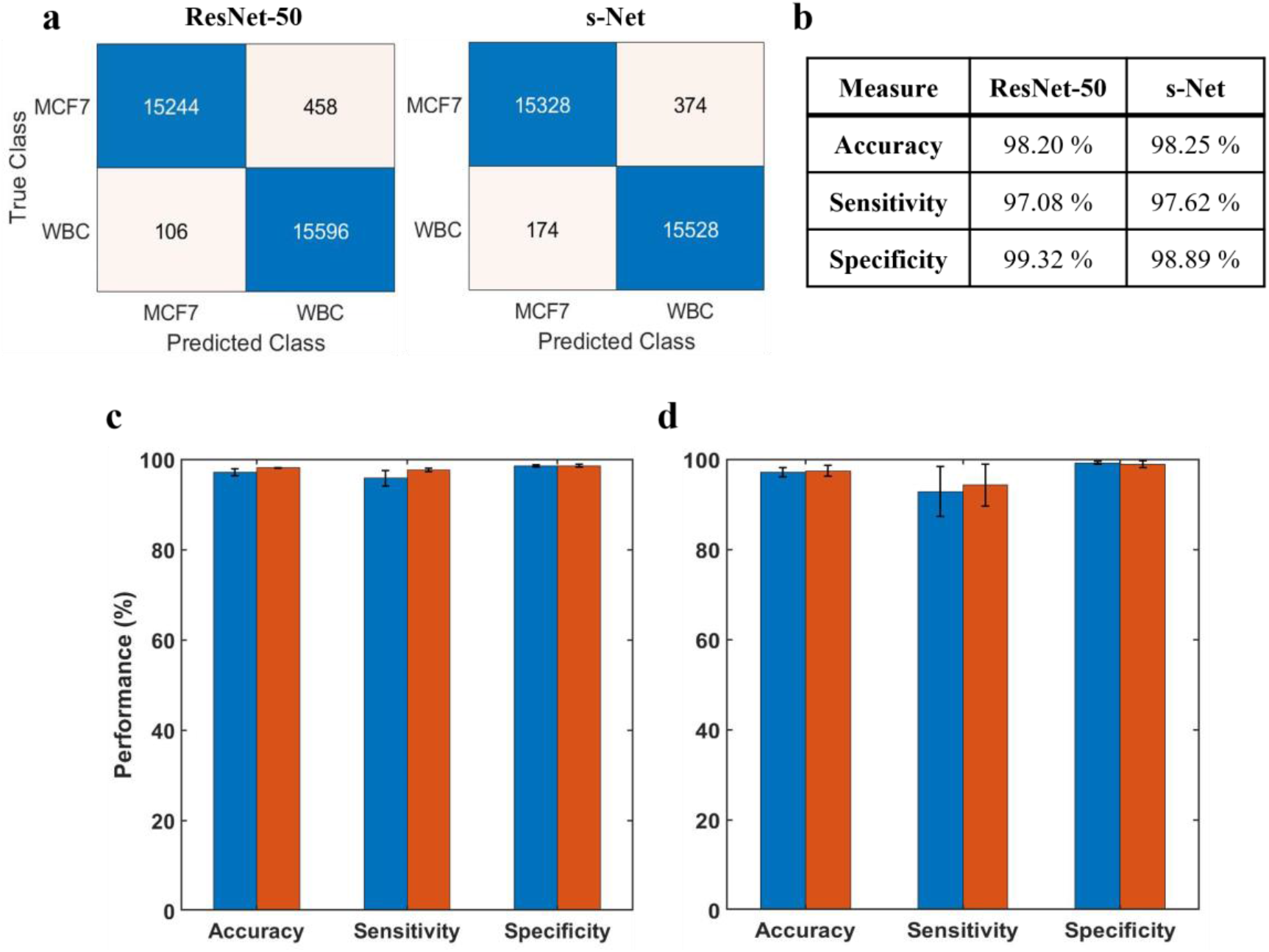
Evaluation of ML model performance. (a) Confusion matrices showing number of TPs, TNs, FPs and FNs for both ResNet-50 and s-Net from a representative trial (b) ML performance is evaluated by determining the accuracy, sensitivity and specificity computed on the test dataset. Results are shown for both network architectures tested (c) Repeatability of the trained CNN. The network is serially trained multiple times with the same dataset; error bars indicate the standard deviation of five trials; (d) Inter-trial variability of the CNN. The trained CNN is tested on multiple datasets; error bars indicate the standard deviation of three independent trials. In (c) and (d), blue corresponds to ResNet-50 and red corresponds to s-Net.

The ML training process is generally subjected to a certain degree of uncertainty introduced by the random initialization of learnable Network weights. This could lead to variations in the classification performance and must be accounted for. To evaluate model repeatability, the model was trained and tested with the same dataset multiple times. Results are reported in Fig. 4c, where the height of the bar indicates the mean value of 5 unique trials. From the standard deviations, we infer that the variability in network performance is inconsequential. Additionally, we evaluated the robustness of the trained model by testing it with datasets generated from 3 different trials (Fig. 4d). In this case, the model was trained only once prior to testing. Once again, we observe very low variability between the trials.

It is evident that the performance of the s-Net is on par with the deeper, over parametrized ResNet-50 network. This comparable performance, in conjunction with its fast computation time make it a suitable candidate to be used for the enumeration task. We use s-Net in subsequent studies.

### 2.4 Detection of tumor cells from artificially mixed datasets using decision thresholding

In spite of the s-Net’s encouraging performance, its application to spiked samples is hindered by the presence of excessive false positives. By calculating the mean from five trials, we find that the FPR is 0.014, which indicates that the s-Net misclassifies approximately 14 WBCs out of 1000 as MCF-7 cells. This is deemed to be too high for our task and results in a significant overestimation of target cell counts. To understand this in the context of our work, let us consider the following scenario: in our trials, as low as 10 MCF-7 cells need to be accurately enumerated from a high background of ~50,000 WBCs. At the current FPR, our Network would misclassify ~700 WBCs as MCF-7s which is clearly not acceptable since from a clinical perspective it would result in a significant confounding of true CTC counts.

In this section, we discuss decision gating, a strategy to reduce the false positive rate associated with classification ^50^. This method involves the application of a strict threshold on the output probabilities generated by the ML Network in order to classify cells. The schematic in Fig. 5a gives an overview of the process. First, mixed population datasets are generated *in silico* from pure population PoBF images of MCF-7 cells and WBCs. For every image fed to the s-Net in a serial fashion, the Softmax layer outputs class probabilities: *P_MCF7_* and *P_WBC_*. A cell is classified as MCF-7 if *P_MCF−7_* > 0.5. To decrease the FPR, it is necessary to use a decision threshold *α* greater than 0.5.

**Figure 5.**
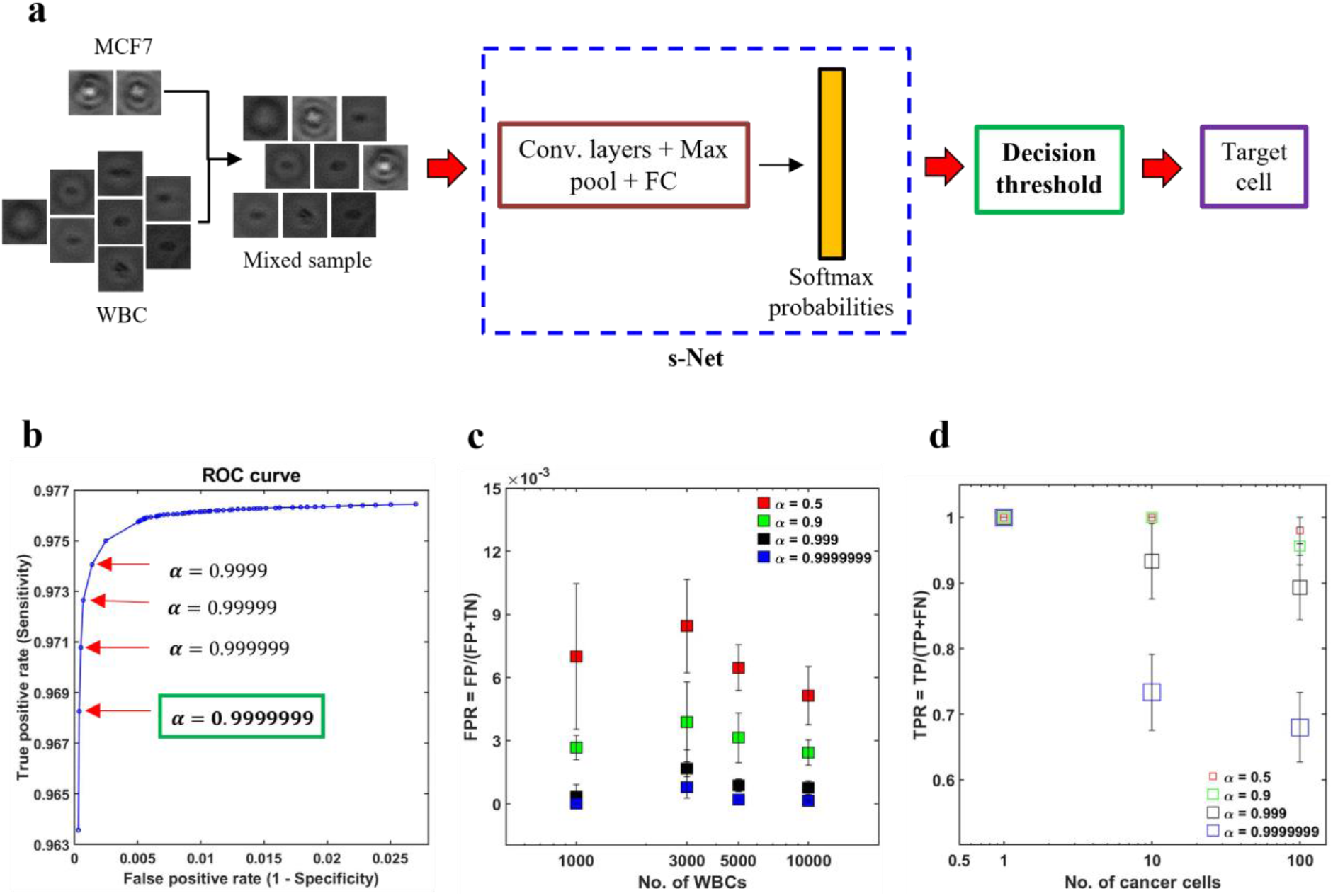
Reducing the false positives (FPs) using decision thresholding. (a) Schematic illustrating the application of decision thresholding for cancer cell classification; (b) Selection of optimal threshold using receiver operator characteristic (ROC) analysis. A stringent threshold (*α*) of 0.9999999 is selected as it yielded the minimum FPR without any further change; (c) To demonstrate the applicability of the selected threshold in reducing the FPR, four artificially mixed samples were generated *in silico*. In these datasets, the number of MCF-7 cancer cells (*N_MCF−7_*) was kept constant at 100, while the number of WBCs (*N_WBC_*) was varied from 1,000-10,000. It was ensured that no overlap between the two cell types existed across the four datasets. For increasing WBC counts, the FPR is plotted for 4 values of *α*: 0.5 (red), 0.9 (green), 0.999 (black) and 0.9999999 (blue). Error bars indicate the standard deviation of three trials (n=3); (d) Effect of *α* on the TPR is shown. In these trials, three datasets were generated, wherein, *N_WBC_* was fixed at 5,000 and *N_MCF−7_* was varied from 1-100. Error bars indicate the standard deviation of three trials (n=3).

To determine the optimal value of *α* we conducted receiver operator characteristic (ROC) analysis, where model sensitivity (Se) and specificity (Sp) are computed on the testing set for different values of *α*. The trained s-Net and test dataset used is the same as that discussed in the previous section. The ROC curve shown in Fig. 5b shows the effect of varying the decision threshold on the true positive and false positive rates. Generally, an increase in decision threshold value *α* results in a reduction of both TPR and FPR. The lowest FPR was obtained at *α* = 0.9999999 which was selected as the optimal threshold. Higher values did not result in any further change in FPR.

The next step is to test the applicability of the selected decision threshold by testing it on artificially mixed datasets containing MCF-7 cells and WBCs. First, we sought to evaluate the effect of *α* on the FPR. In order to do this, four independent datasets were created where the number of MCF-7 cells (*N_MCF−7_*) was fixed at 100 but the number of WBCs (*N_WBC_*) was made to range from 1,000 – 10,000. The rationale for doing so was that the FPR depends solely on the classification statistics computed on the negative class, i.e., WBCs in our case. For this reason, *N_MCF−7_* was kept constant. For each WBC count tested, we plot the FPR at four different decision thresholds. As shown in Fig. 5c, increasing the decision thresholding helps to significantly reduce the false positive rate. At the selected threshold, i.e., *α* = 0.9999999, the mean FPR is measured to be 2.77 × 10^−4^. Additionally, contrary to the case of low *α*, where FPR is much more sensitive to the number of WBCs needed to be screened in the sample, at the selected threshold we find that the FPR is not varying significantly with WBC counts. This invariance in FPR suggests that as the number of background WBCs in the sample (*N_WBC_*) increases, the number of false positives also increase.

Undoubtedly, while the thresholding strategy has been effective in lowering the FPR, imposing such a strict boundary invariably lowers the number of true positives. This makes it important to evaluate the extent of decrease in TPR and assess whether this is acceptable or not. To achieve this objective, we created three mixed datasets where only the number of cancer cells were varied: *N_MCF−7_* = 1,10,100 while the number of WBCs was kept constant (*N_WBC_* = 5000). We find that TPR depends on both the number of MCF-7s as well as *α* (Fig. 5d). Evaluating at *α* = 0.9999999, we find that TPR decreased to about 0.7 for *N_MCF−7_* = 10 and 100 respectively. In the dataset with only a single cancer cell, our s-Net model is able to detect it robustly at the selected threshold.

Overall, our analysis suggests that both the FPs and TPs depend on the decision threshold as well as counts of WBCs and MCF-7s. This highlights the potential limitation of the deep-learning model for label-free CTC enumeration. CTCs detected as WBCs would result in underestimating the tumor burden which could prove to be fatal for the cancer patient. On the other hand, scoring WBCs as CTCs would result in overpredicting the actual CTC counts, with the possibility of subjecting patients to unnecessary treatment regimens. Acknowledging this limitation, we wondered how well the DHM-ML approach would characterize varying loads of MCF-7s in actual blood samples, which we discuss in the next section.

### 2.5 DHM-ML-assisted enumeration of tumor cells spiked into lysed blood

We applied the decision thresholding method to enumerate tumor cells from actual mixed samples containing a background of WBCs. Briefly, fluorescently labelled MCF-7 breast cancer cells were spiked at different target concentrations: 0, 10, 100 and 1000/mL. The concentration of WBCs was kept constant at 90,000/mL – this high concentration was chosen as it might present a worst-case scenario for ML detection. To achieve flow-based interrogation of tumor cells, 10100 holograms of mixed samples containing MCF-7 cells and WBCs were captured to image a total sample volume of 1 mL. The total number of cells screened ranged from 35,000 - 50,000. Automated ML-based enumeration was carried out according to the earlier-mentioned procedure (Fig. 2a). For validating the detected concentrations, we rely not on the theoretical target counts but obtain separate ground truth counts. This was done by collecting the entire DHM-imaged sample from the outlet of the sheath device (~1.4 ml total volume from both sample and sheath fluid) and performing fluorescence imaging to obtain the actual counts of MCF-7 cells (details in Sec. 5.7). Establishing a robust validation strategy was important since sample handling can introduce variability and serial dilution could lead to high errors at extremely low spiked concentrations.

Fig. 6a shows the enumeration results of spiked tumor cells as predicted by the s-Net model with the data plotted on a log-log scale. To improve the accuracy of the recovered target cells, the optimized decision threshold *α* = 0.9999999 was applied. We find that the enumerated MCF-7 concentrations increase with increasing spiked cell concentrations, although the actual numbers do not quantitatively agree (Fig. 6b). Thus, the DHM-ML approach can sufficiently distinguish between varying loads of cancer cells at high WBC background counts indicating that it can be considered as a qualitative screening tool to identify blood samples that have high number of CTCs.

**Figure 6.**
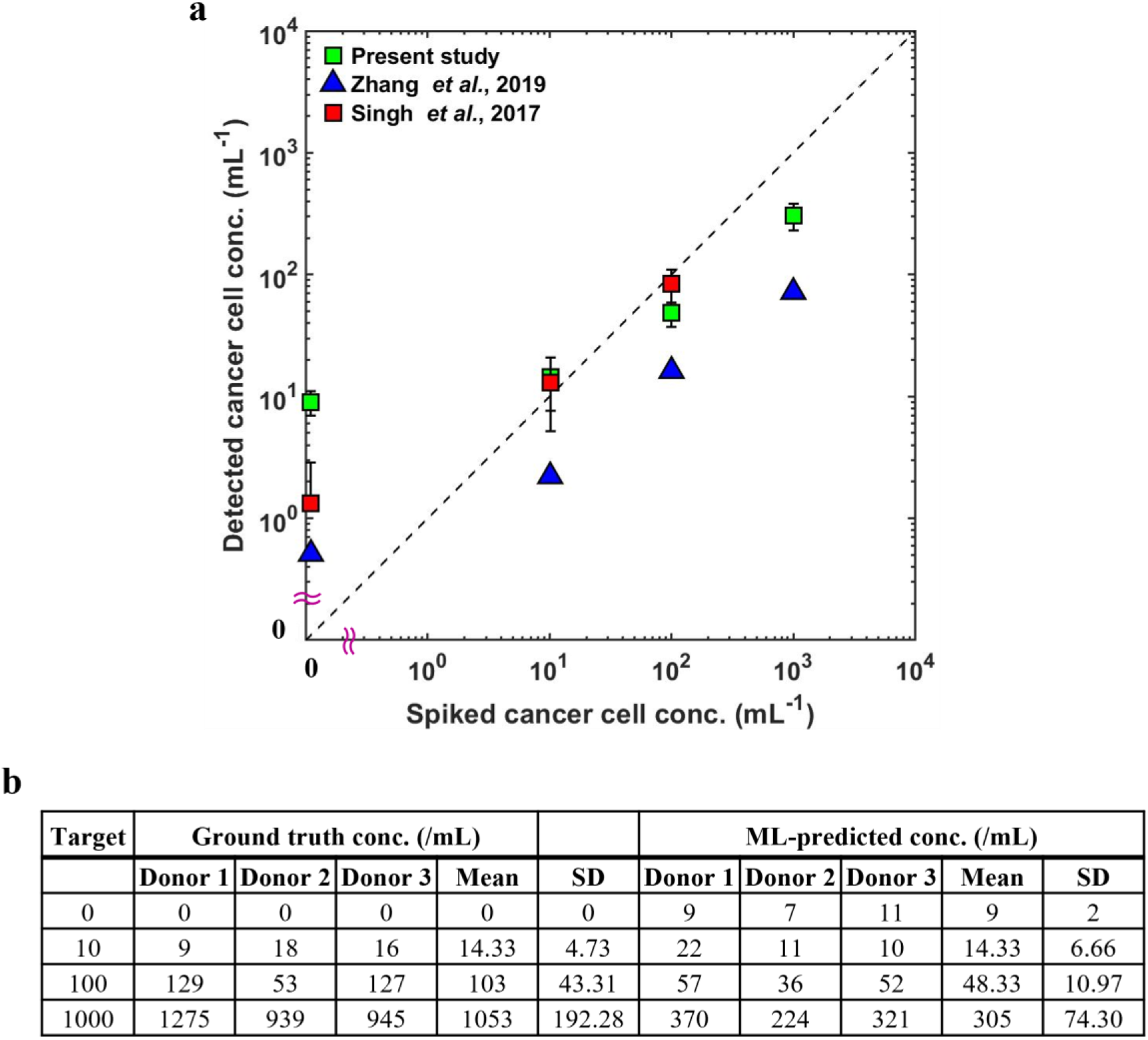
ML-based detection and enumeration of MC F7 breast cancer cells spiked into lysed blood at various concentrations. (a) Enumeration of MCF7 breast cancer cells spiked at various concentrations into lysed blood, background WBC concentration is 90,000 per mL. The abscissa shows the target counts of cancer cells spiked into lysed blood. Error bars denote the standard deviations across three independent trials performed. Results from the study by Zhang *et al.* as well as our laboratory’s previous work (Singh *et al.*) are also overlaid in the same plot (latter study did not test C=1000/mL). Data in blue represents the mean concentration of three separate trials. The analytical limit of detection (LoD) as determined from the negative control data is 13 per mL; (b) Table shows the detected MCF-7 concentrations as predicted by ML along with the corresponding fluorescent ground truth counts. Mean and standard deviations of 3 trials conducted using independent donor blood specimens are also reported.

As part of establishing the screening or enumeration potential of an assay, it is critical to report the limit of detection (LoD). This is defined as the lowest target cell concentration that can be reliably detected by an assay and is given by: *LoD* = *μ_NC_* + 2 *σ_NC_* ^51^. Here, *μ_NC_*, *σ_NC_* denote mean and standard deviation of the negative control results, i.e., the sample does not contain any analyte. To calculate the LoD for our system, we performed 3 negative control trials, where the samples contained WBCs but no MCF-7 cells (Fig. 6b). This yielded an LoD of 13 target cells/mL sample analyzed (*μ_NC_* = 9, *σ_NC_* = 2), which is comparable to typical values reported in the literature ^34,50^.

### 2.6 Comparison of present work with literature studies

It is useful to compare the results of our work with those from literature where holographic imaging was used to distinguish cancer cells and WBCs. Zhang *et al.*^50^ enumerated MCF-7 breast cancer cells spiked into diluted blood samples using lens-less holographic imaging and a densely connected deep neural network (P3D-CNN). The authors initially enriched their sample using labelled magnetic beads that selectively bind to MCF-7 cells. Since the sample still contained a high background of 1.6 × 10^6^ blood cells/mL, the bead-labelled MCF-7 cells were actuated under a deterministic, periodic magnetic force. Upon actuation, the bead-bound target cells exhibited different spatio-temporal fingerprints which was used to improve selectivity. Briefly, the recorded inline holographic images were first subjected to a computational drift correction step to account for the field-induced fluid motion in the imaging tube. This was followed by numerical reconstruction, periodic motion estimation and 3D localization. Finally, the in-focus amplitude and phase images were fed to a densely connected deep neural network (P3D-CNN) which generated Softmax probabilities enabling classification of target cells using a stringent decision threshold (*α* = 0.999999) on the output probabilities.

As shown in Fig. 6a, the data of Zhang *et al*.^50^ lies lower than the anticipated detected concentrations, indicating that our DHM-ML approach performs better even though the decision thresholds are similar. Moreover, they used magnetic beads to perform enrichment, which only selects cells positive for the labelled beads unlike our label-free approach. In addition, the flow-through approach that we demonstrated has the potential to process larger sample volumes compared to the static sample visualization they performed. Finally, we note that Zhang *et al*. ^50^ used a lowest target cell concentration of 10 cells/mL and did not report an LoD based on testing three negative control samples, similar to what we did in our study.

In another study, Singh *et al.*^34^ used a decision tree classification strategy based on the CART algorithm to enumerate two MCF-7 breast cancer cells spiked into lysed blood in flow using inline holographic images. Their detected cell concentration is shown in Fig. 6a and is comparable to our data with an LoD that is lower: 4.33 cells/mL. Also, the number of background cells of 450,000 WBCs per sample was higher. Compared to the work of Singh *et al.*^34^, this work has made three advances: (1) our portable holographic setup can be easily integrated with standard inverted microscopes, which only requires the 3D-printed housing to be mounted on the microscope stage. This is in contrast to the elaborate optical setup used by Singh *et al.*^34^. The 3D-printed portable laser torch design allows easy translation to research settings (2) Unlike Singh *et al.*^34^, we used a sheath-flow device which helped to eliminate fringes from channel side walls, and reduce the presence of slow-moving near-wall cells. This wall-less approach improves the accuracy of cell localization and enumeration. (2) our approach of using CNN allows image-based classification without any intermediate feature extraction and enables extraction of the best set of discriminators for the classification task^52,53^. This is advantageous because one need not worry about the best set of features to be used which requires domain expertise especially when ML models for different types of cancer cells (e.g., prostate, lung cancer etc..) need to be developed.

## 3. Discussion

Immunostaining is the current gold-standard for identifying CTCs from a background of blood cells. Despite being highly selective, this approach is time-consuming requiring multiple processing steps, and is dependent on the quality of antibodies. Importantly, this approach limits some critical CTC applications like ex vivo culture expansion of CTCs, and omic analysis where there are opposing needs to access live CTCs but also in selecting patients with adequate CTC counts. This gap can be addressed by developing staining-free approaches to detect CTCs, and our effort is among the first few studies that have attempted to tackle this problem using deep learning models. Our results show that the DHM-ML strategy is promising, but additional research efforts are needed to translate this approach to blood samples of cancer patients, which we discuss below.

### 3.1 Integration of CTC enrichment technologies can increase accuracy of ML detection

In our DHM-ML study we used a sheath device in which a mixture of MCF-7 cells and WBCs were evaluated. Many CTC isolation technologies enrich the CTC population using mechanical filters^54,55^, inertial focusing^8,22,56,57^ and negative selection^58–60^. Our approach is compatible with any of these approaches since the enriched samples can be collected and introduced into the sheath device for enumeration. This a major advantage of our DHM-ML strategy as it allows broader application to a variety of CTC-enrichment technologies.

Our results show that when the background WBC counts are high, the ML model needs to balance between FPR and TPR, but at low WBC counts it can perform with greater accuracy. Specifically, for *N_WBC_* = 1000, our ML model predicts 0 false positives (for *α* = 0.9999999) which is quite promising. Given the limitation of the deep-learning approach for high WBC counts, it is important to apply enrichment strategies to reduce the background WBC counts, as this will lead to lower FPs. Different CTC isolation technologies report different levels of sample purity after enrichment, *i.e*., producing different background WBC counts. For example, label-free technologies such as the Vortex HT chip^29^ and Labyrinth^30^ produce background WBC counts in the range of 25 – 1000 per mL sample. At such high levels of purity, we anticipate that the LoD can be lowered and the FPR reduced even further. Therefore, implementing such enrichment strategies prior to DHM-ML detection should bring a major advance in staining-free CTC enumeration.

### 3.2 Expanding pDHM-ML approach to patient samples requires dedicated computational hardware

Access to live CTCs is the next frontier in liquid-biopsy research. However, significant intrapatient and inter-patient variability exists in CTC counts^61^. Some of the blood samples may not even contain CTCs or may have very low counts bringing ambiguity into the type of downstream assays that could be pursued. Therefore, there is a need for a quick screening tool that can assess live CTCs and our DHM-ML technique presents a promising avenue to address this gap. Given that CTCs are rare, high volume of blood needs to be processed which may increase the computational burden on hologram processing. However, implementing an enrichment step eliminates the need to computationally enumerate blood cells. As an example, the Labyrinth chip is able to process 5 mL of patient whole blood and deliver a 2 mL CTC-enriched sample volume. Using our pDHM setup, this sample can be flowed through our sheath channel and the entire sample can be imaged by recording 21,000 experimental holograms. It takes ~5 seconds to process a single hologram on a standard desktop computer with 4 CPU cores. At this rate, we estimate that the entire sample can be analyzed, and tumor cell counts enumerated from 5 mL whole blood in ~29 hours. This computational time can be reduced by integrating dedicated hardware such as Tensor core-enabled GPUs ^62^ or increasing the number of CPU cores ^63^ for parallelized operations to fast-track our analysis.

### 3.3 Developing ML models with cancer patient blood samples

In our work, we used MCF-7 cells as a proxy for CTC, which is a limitation for translating our results to patient-derived CTCs. The challenge with developing ML models with real CTCs is generation of ground-truth data. Fluorescent markers capable of tagging live CTCs, coupled with additional morphological features (e.g., cell size, nucleus-to-cytoplasm ratio) could be used to label CTCs for ML model development. A few studies are emerging addressing this gap. For example, Wang *et al.*^28^ used a Carbonic Anhydrase 9 antibody in conjunction with Calcein AM to differentiate CTCs from blood cells. In another study by Shao *et al.*^64^, CTCs from prostate cancer patients were successfully labeled using a class of near-infrared Heptamethine Carbocyanine dyes. Although these advances are promising, further work is needed to develop robust markers capable of labeling CTCs across different types of cancers.

Another major hurdle for ML model development is that since these CTCs occur at extremely low frequencies (1-100 cells/mL) in blood, collecting an adequate amount of ground truth data would require analysis of many blood specimens. Collecting large number of blood samples can be a logistical challenge. There are two main approaches to deal with this challenge: (1) recently, DL-based approaches such as Generative Adversarial Networks (GANs) have demonstrated sufficiently good performance with small or unbalanced datasets and can be explored; (2) for generating larger training datasets, a combination of patient-derived CTCs and *in vitro* cell lines may be used for the positive class^28^.

## 4. Conclusion

In this work, we demonstrated staining-free enumeration of tumor cells in lysed blood samples using digital holographic microscopic imaging and machine learning. A custom-designed 3D printed laser torch was engineered to simplify the holography microscopy setup. A sheath-flow device was used to improve the accuracy of cell detection and enumeration. Numerical reconstruction was implemented to obtain the plane of best focus images which were used to train and develop deep-learning models.

First the classification performance was evaluated by contrasting two deep-learning network architectures: (1) ResNet-50 and (2) a custom s-Net. We found that the performance of the shallow s-Net was comparable to the ResNet and was selected as the network of choice due to its simplicity and reduced computational burden. Model repeatability was evaluated by training and testing the model on the same dataset multiple times. Furthermore, to evaluate the robustness of the trained s-Net, we tested it on datasets generated from three separate trials (n=3). In both cases, s-Net demonstrated good generalization capability.

To apply the trained model to mixed samples, it was deemed necessary to significantly lower the FPR. This was achieved using decision thresholding applied on the Softmax probabilities predicted by the network. After determining the optimal threshold, we tested its applicability on mixed samples generated *in silico*. At a threshold of 0.9999999, the FPR was found to decrease by almost two orders of magnitude, thereby greatly reducing the number of misclassified WBCs. Finally, the decision-thresholding-enabled CNN approach was tested on actual mixed samples containing MCF-7 breast cancer cells spiked at various target concentrations into lysed blood. In general, we achieved good enumeration performance at all the tested concentrations with an LoD of 13 target cells/mL.

In summary, our DHM-ML strategy represents the first step in developing CNN-based deep learning-based frameworks for label-free enumeration of tumor cells among a background of WBCs. While transitioning to patient samples still requires advances, the approach presented holds promise to be applied to other types of cancer cells, potentially lending to a quick screening tool for label-free enumerations of CTCs.

## 5. Materials and Methods

### 5.1 Cell culture

A stock of MCF-7 breast cancer cells was obtained from American Type Cell Collection (ATCC, Manassas, VA). The cells were cultured using Dulbecco’s modified Eagle medium (DMEM, Gibco, Gaithersburg, MD) supplemented with 10% Fetal Bovine Serum (FBS, Gibco), 1% Penicillin/Streptomycin (Gibco) and 1% sodium pyruvate (Gibco). They were incubated at 37°C in a humidified atmosphere containing 5% CO_2_ to maintain ambient conditions.

### 5.2 Fluorescent labeling of tumor cells

MCF-7 Cells were fluorescently tagged with CellTracker™ Green CMTPX dye (Invitrogen™, Waltham, MA). A stock solution of 10 mM was prepared according to manufacturer’s instructions. From this, a 10 *μM* working solution was prepared by diluting the stock in serum-free media. This solution was then added to the adherent cells in the tissue culture flask followed by an incubation step at 37 °C for 45 minutes. Subsequently, the cells were washed thrice with 1X phosphate buffered saline (PBS, Gibco) to remove any excess dye. The washed cells were trypsinized and resuspended in 500 *μL* of filtered, fresh media.

### 5.3 Isolation of WBCs

Fresh human whole blood was purchased from BioIVT (Westbury, NY). 10 mL of blood was drawn from healthy donors into vacutainer tubes containing K2EDTA as anticoagulant and shipped on the same day. WBCs were isolated using ACK lysis buffer (Quality Biological, Gaithersburg, MD). First, 1 mL whole blood was incubated with 10 mL lysis buffer at room temperature for 5 minutes. This was followed by centrifugation at 2000 rpm for 5 minutes. After discarding the supernatant, the pellet was resuspended and mixed gently in 5mL of lysis buffer following which the incubation and spinning steps were repeated. Subsequently, the supernatant was extracted and care was taken to ensure that the red RBC pellet was removed without disturbing the yellowish-white WBC pellet that was finally resuspended in 1X PBS.

### 5.4 Sample preparation

MCF-7 cells and WBC stock suspensions were passed through a 30 *μm* filter (CellTrics, Norderstedt, Germany) to remove any debris or other impurities. Cell concentrations were measured using a Neubauer hemocytometer. MCF-7 cells were counted three times and the average was recorded as the stock concentration. Pure population samples contained a concentration of 100,000 cells/mL. Mixed samples containing MCF-7 cells at spiked concentrations of 10, 100 and 1000 /mL and WBCs (90,000 per mL) were prepared in 1X PBS in accordance with Table S1.

### 5.5 Microfabrication

The sheath microchannel, 800 *μm* wide and 330 *μm* deep was fabricated using soft lithography ^65^. Negative photomasks were first designed in AUTOCAD (v. 2019, Autodesk, San Rafael, CA) and printed. SU-8 2050 (MicroChem Corporation, Westborough, MA), a negative photoresist, was used to prepare the mold. To achieve the large depth, a 2-step spin coating procedure was followed. At each step, the target height was selected to be 170 *μm*. The wafer with the etched pattern underwent trichloro (1H, 1H, 2H, 2H-perfluorooctyl) silane (Sigma Aldrich, St. Louis, MO) treatment for 24 hours. To prepare the devices, PDMS pre-polymer and the curing agent (Sylgard 184 Silicone Elastomer kit, Dow Corning Inc., Midland, MI) were mixed in a 10:1 w/w ratio, degassed, poured on the mold and cured in a 70 °C oven for 2 hours. The cured PDMS replicas were cut and peeled. Inlet and outlet fluidic ports were made using a 1 mm hole puncher (Instech Laboratories, Plymouth Meeting, PA). This was followed by plasma treatment (Harrick Plasma, Ithaca, NY) for 90 seconds to irreversibly bond the PDMS cutout to a 25 mm x 75 mm x 1 mm glass slide (Thermo Fisher, Waltham, MA). The hydrophilic devices were then incubated at 70 °C for 15 minutes to strengthen the surface bonding. Prior to experiments, 1 mL of 1% (w/v) Pluronic® F-127 (Sigma Aldrich, St. Louis, MO) prepared in 1X PBS was flowed through the microchannel at 100 *μL/min* ^30^. This was done to prevent cell adhesion to the PDMS channel walls.

### 5.6 Digital holographic microscopy (DHM) imaging

The holographic setup consists of a laser torch (LDM 635, Thorlabs, NJ, USA) placed in a 3D-printed housing. The emitted laser beam (*λ* = 635 *nm*, 4 mW, diameter: 3 mm x 5 mm) serves as a coherent light source and is operated in the continuous wave (CW) mode. The imaging field of view (FOV) is 800 x 800 (in pixels) and the depth of field (or channel depth) is 330 *μm*. The flow is actuated by a syringe pump (PHD 2000, Harvard Apparatus). The flow rates of the sample and sheath fluid streams are 2.5 and 0.5 mL/min respectively, resulting in a total flow rate of 3.5 mL/min. A frame rate of 420 frames/sec is used in our experiments. The holograms are magnified by a 20X (1*μm/pix*.) objective (20X, NA = 0.45, Olympus) with the hologram plane located 200 *μm* below the microchannel floor. Subsequently, they are recorded on a CMOS sensor of a high-speed camera (Phantom v310, Vision Research), facilitated by the PCC software (Phantom, Vision Research). An exposure time of 35 *μs* is used. 10,100 raw holograms are captured in order to image 1 mL of sample volume which takes about 24 seconds at the imposed frame rate.

### 5.7 Post-DHM fluorescence imaging for ground truth count generation

The DHM-imaged sample was collected and transferred to a 96-well plate (Greiner Bio-One, Monroe, NC), with each well containing 150 *μl* of sample volume. After allowing the cells to settle for about 20 minutes, imaging was performed using an Olympus IX81 epifluorescence microscope (Massachusetts, USA). By using a programmable stage (Thorlabs, New Jersey, USA), images were recorded in an automated fashion using the Slidebook 6.1 software (3i Intelligent Imaging Innovations Inc., Denver, USA). A digital monochrome camera (Hamamatsu, ImagEM X2 EM-CCD, New Jersey, USA) was used for capture. Fluorescence FITC images were acquired at 20X (512×512 pixels, 0.8 *μm/pix*.) objective magnification with an exposure time of 100 ms. After acquisition, ground truth counts of the FITC-positive MCF-7 cells were obtained from the images. It is important to note that WBCs were not tagged fluorescently and therefore are not present in these images.

### 5.8 Computational analysis

The automated analysis involving processing of raw holograms and training/testing of the ML models was performed on a desktop computer (Intel(R) Core(TM) i7-7700 CPU, 3.60GHz, 16 GB RAM) using MATLAB (R2021b; MathWorks Inc., Natick, Massachusetts) software. The average processing speed per hologram is 5 seconds. For training the deep ResNet-50 Network, a GPU-enabled (NVIDIA GeForce GTX 1060, 6GB) system (AMD Ryzen Threadripper 2990WX 32-Core Processor, 3.00 GHz, 32 GB RAM) was used. Model training hyperparameters such as number of convolutional filters, filter size, number of epochs, minibatch size and learning rate of the optimizer were tuned to achieve optimal generalized classification performance (details in Sec. 2.3).

## 6. Author contributions

AG and SAV conceptualized the study, AG designed and conducted the experiments, developed the analysis framework and generated all the data. AG and SAV wrote the paper. SAV and HSS provided feedback on the manuscript and supervised the project.

## 7. Acknowledgements

This work was funded by the Cancer Prevention Research Institute of Texas (CPRIT, Grant no. RP190658). We are grateful to Siddhartha Gupta and Kevin Loftis for the design and testing of the 3D-printed laser torch.

## References

1. Lin Z, Luo G, Du W, Kong T, Liu C, Liu Z. Recent Advances in Microfluidic Platforms Applied in Cancer Metastasis: Circulating Tumor Cells’ (CTCs) Isolation and Tumor-On-A-Chip. Small 2020; 16(9): e1903899.

2. Cristofanilli M, Budd GT, Ellis MJ, et al. Circulating tumor cells, disease progression, and survival in metastatic breast cancer. N Engl J Med 2004; 351(8): 781–91.

3. Hayes DF, Cristofanilli M, Budd GT, et al. Circulating tumor cells at each follow-up time point during therapy of metastatic breast cancer patients predict progression-free and overall survival. Clinical Cancer Research 2006; 12(14): 4218–24.

4. Nagrath S, Sequist LV, Maheswaran S, et al. Isolation of rare circulating tumour cells in cancer patients by microchip technology. Nature 2007; 450(7173): 1235–9.

5. Mali SB, Dahivelkar S. Liquid biopsy = Individualized cancer management: Diagnosis, monitoring treatment and checking recurrence and metastasis. Oral Oncol 2021; 123.

6. Burinaru TA, Avram M, Avram A, et al. Detection of Circulating Tumor Cells Using Microfluidics. Acs Comb Sci 2018; 20(3): 107–26.

7. Hong B, Zu Y. Detecting circulating tumor cells: current challenges and new trends. Theranostics 2013; 3(6): 377–94.

8. Hou HW, Warkiani ME, Khoo BL, et al. Isolation and retrieval of circulating tumor cells using centrifugal forces. Sci Rep 2013; 3: 1259.

9. Mashhadi SMY, Kazemimanesh M, Arashkia A, et al. Shedding light on the EpCAM: An overview. J Cell Physiol 2019; 234(8): 12569–80.

10. Sieuwerts AM, Kraan J, Bolt J, et al. Anti-Epithelial Cell Adhesion Molecule Antibodies and the Detection of Circulating Normal-Like Breast Tumor Cells. J Natl Cancer I 2009; 101(1): 61–6.

11. Stott SL, Hsu CH, Tsukrov DI, et al. Isolation of circulating tumor cells using a microvortexgenerating herringbone-chip. Proc Natl Acad Sci U S A 2010; 107(43): 18392–7.

12. Thiery JP, Lim CT. Tumor Dissemination: An EMT Affair. Cancer Cell 2013; 23(3): 272–3.

13. Hyun K-A, Kwon K, Han H, Kim S-I, Jung H-I. Microfluidic flow fractionation device for label-free isolation of circulating tumor cells (CTCs) from breast cancer patients. Biosensors and Bioelectronics 2013; 40(1): 206–12.

14. Tan SJ, Lakshmi RL, Chen P, Lim W-T, Yobas L, Lim CT. Versatile label free biochip for the detection of circulating tumor cells from peripheral blood in cancer patients. Biosensors and Bioelectronics 2010; 26(4): 1701–5.

15. Warkiani ME, Guan G, Luan KB, et al. Slanted spiral microfluidics for the ultra-fast, label-free isolation of circulating tumor cells. Lab on a Chip 2014; 14(1): 128–37.

16. Hur SC, Henderson-MacLennan NK, McCabe ER, Di Carlo D. Deformability-based cell classification and enrichment using inertial microfluidics. Lab on a Chip 2011; 11(5): 912–20.

17. Moon H-S, Kwon K, Kim S-I, et al. Continuous separation of breast cancer cells from blood samples using multi-orifice flow fractionation (MOFF) and dielectrophoresis (DEP). Lab on a Chip 2011; 11(6): 1118–25.

18. Huang S-B, Wu M-H, Lin Y-H, et al. High-purity and label-free isolation of circulating tumor cells (CTCs) in a microfluidic platform by using optically-induced-dielectrophoretic (ODEP) force. Lab on a Chip 2013; 13(7): 1371–83.

19. Gertler R, Rosenberg R, Fuehrer K, Dahm M, Nekarda H, Siewert JR. Detection of circulating tumor cells in blood using an optimized density gradient centrifugation. Molecular Staging of Cancer: Springer; 2003: 149–55.

20. Harb W, Fan A, Tran T, et al. Mutational Analysis of Circulating Tumor Cells Using a Novel Microfluidic Collection Device and qPCR Assay. Transl Oncol 2013; 6(5): 528-+.

21. Ozkumur E, Shah AM, Ciciliano JC, et al. Inertial focusing for tumor antigen-dependent and-independent sorting of rare circulating tumor cells. Sci Transl Med 2013; 5(179): 179ra47.

22. Warkiani ME, Khoo BL, Wu L, et al. Ultra-fast, label-free isolation of circulating tumor cells from blood using spiral microfluidics. Nat Protoc 2016; 11(1): 134–48.

23. Kapeleris J, Kulasinghe A, Warkiani ME, et al. Ex vivo culture of circulating tumour cells derived from non-small cell lung cancer. Transl Lung Cancer Res 2020; 9(5): 1795–809.

24. Maheswaran S, Haber DA. Ex Vivo Culture of CTCs: An Emerging Resource to Guide Cancer Therapy. Cancer Res 2015; 75(12): 2411–5.

25. Yu M, Bardia A, Aceto N, et al. Cancer therapy. Ex vivo culture of circulating breast tumor cells for individualized testing of drug susceptibility. Science 2014; 345(6193): 216–20.

26. Wei TT, Zhu DL, Yang Y, Yuan GD, Xie HY, Shen RM. The application of nano-enrichment in CTC detection and the clinical significance of CTCs in non-small cell lung cancer (NSCLC) treatment. Plos One 2019; 14(7).

27. Li JP. Significance of Circulating Tumor Cells in Nonsmall-Cell Lung Cancer Patients: Prognosis, Chemotherapy Efficacy, and Survival. J Healthc Eng 2021; 2021.

28. Wang S, Zhou Y, Qin X, Nair S, Huang X, Liu Y. Label-free detection of rare circulating tumor cells by image analysis and machine learning. Sci Rep 2020; 10(1): 12226.

29. Che J, Yu V, Dhar M, et al. Classification of large circulating tumor cells isolated with ultra-high throughput microfluidic Vortex technology. Oncotarget 2016; 7(11): 12748–60.

30. Lin E, Rivera-Baez L, Fouladdel S, et al. High-Throughput Microfluidic Labyrinth for the Label-free Isolation of Circulating Tumor Cells. Cell Syst 2017; 5(3): 295–304 e4.

31. Drucker A, Teh EM, Kostyleva R, Rayson D, Douglas S, Pinto DM. Comparative performance of different methods for circulating tumor cell enrichment in metastatic breast cancer patients. Plos One 2020; 15(8).

32. Xu L, Mao X, Imrali A, et al. Optimization and Evaluation of a Novel Size Based Circulating Tumor Cell Isolation System. PLoS One 2015; 10(9): e0138032.

33. Dhar M, Pao E, Renier C, et al. Label-free enumeration, collection and downstream cytological and cytogenetic analysis of circulating tumor cells. Scientific Reports 2016; 6.

34. Singh DK, Ahrens CC, Li W, Vanapalli SA. Label-free, high-throughput holographic screening and enumeration of tumor cells in blood. Lab Chip 2017; 17(17): 2920–32.

35. Milgram JH, Li WC. Computational reconstruction of images from holograms. Appl Optics 2002; 41(5): 853–64.

36. Schnars U, Juptner WPO. Digital recording and numerical reconstruction of holograms. Measurement Science and Technology 2002; 13(9): R85–R101.

37. Rappaz B, Marquet P, Cuche E, Emery Y, Depeursinge C, Magistretti PJ. Measurement of the integral refractive index and dynamic cell morphometry of living cells with digital holographic microscopy. Optics Express 2005; 13(23): 9361–73.

38. Kemper B, von Bally G. Digital holographic microscopy for live cell applications and technical inspection. Appl Optics 2008; 47(4): A52–A61.

39. Molaei M, Sheng J. Imaging bacterial 3D motion using digital in-line holographic microscopy and correlation-based de-noising algorithm. Opt Express 2014; 22(26): 32119–37.

40. Choi YS, Lee SJ. Three-dimensional volumetric measurement of red blood cell motion using digital holographic microscopy. Appl Optics 2009; 48(16): 2983–90.

41. Rubin M, Stein O, Turko NA, et al. TOP-GAN: Stain-free cancer cell classification using deep learning with a small training set. Med Image Anal 2019; 57: 176–85.

42. Ugele M, Weniger M, Stanzel M, et al. Label-Free High-Throughput Leukemia Detection by Holographic Microscopy. Adv Sci (Weinh) 2018; 5(12): 1800761.

43. Watanabe E, Hoshiba T, Javidi B. High-precision microscopic phase imaging without phase unwrapping for cancer cell identification. Optics Letters 2013; 38(8): 1319–21.

44. Merola F, Memmolo P, Miccio L, et al. Tomographic flow cytometry by digital holography. Light-Sci Appl 2017; 6.

45. Singh DK, Ahrens CC, Li W, Vanapalli SA. Label-free fingerprinting of tumor cells in bulk flow using inline digital holographic microscopy. Biomed Opt Express 2017; 8(2): 536–54.

46. Memmolo P, Miccio L, Merola F, Gennari O, Netti PA, Ferraro P. 3D morphometry of red blood cells by digital holography. Cytometry A 2014; 85(12): 1030–6.

47. Oe K, Nomura T. Twin-image reduction method using a diffuser for phase imaging in-line digits holography. Appl Optics 2018; 57(20): 5652–6.

48. Shangraw M, Ling HJ. Separating twin images in digital holographic microscopy using weak scatterers. Appl Optics 2021; 60(3): 626–34.

49. Ling HJ, Katz J. Separating twin images and locating the center of a microparticle in dense suspensions using correlations among reconstructed fields of two parallel holograms. Appl Optics 2014; 53(27): G1–G11.

50. Zhang Y, Ouyang M, Ray A, et al. Computational cytometer based on magnetically modulated coherent imaging and deep learning. Light Sci Appl 2019; 8: 91.

51. Armbruster DA, Pry T. Limit of blank, limit of detection and limit of quantitation. Clin Biochem Rev 2008; 29 Suppl 1: S49–52.

52. LeCun Y, Bengio Y, Hinton G. Deep learning. Nature 2015; 521(7553): 436–44.

53. Ren SQ, He KM, Girshick R, Sun J. Faster R-CNN: Towards Real-Time Object Detection with Region Proposal Networks. Adv Neur In 2015; 28.

54. Lim LS, Hu M, Huang MC, et al. Microsieve lab-chip device for rapid enumeration and fluorescence in situ hybridization of circulating tumor cells. Lab on a Chip 2012; 12(21): 4388–96.

55. Vona G, Sabile A, Louha M, et al. Isolation by size of epithelial tumor cells - A new method for the immunomorphological and molecular characterization of circulating tumor cells. American Journal of Pathology 2000; 156(1): 57–63.

56. Warkiani ME, Guan G, Luan KB, et al. Slanted spiral microfluidics for the ultra-fast, label-free isolation of circulating tumor cells. Lab Chip 2014; 14(1): 128–37.

57. Warkiani ME, Khoo BL, Tan DS, et al. An ultra-high-throughput spiral microfluidic biochip for the enrichment of circulating tumor cells. Analyst 2014; 139(13): 3245–55.

58. Mishra A, Dubash TD, Edd JF, et al. Ultrahigh-throughput magnetic sorting of large blood volumes for epitope-agnostic isolation of circulating tumor cells. P Natl Acad Sci USA 2020; 117(29): 16839–47.

59. Lee TY, Hyun KA, Kim SI, Jung HI. An integrated microfluidic chip for one-step isolation of circulating tumor cells. Sensor Actuat B-Chem 2017; 238: 1144–50.

60. Hyun KA, Lee TY, Lee SH, Jung HI. Two-stage microfluidic chip for selective isolation of circulating tumor cells (CTCs). Biosensors & Bioelectronics 2015; 67: 86–92.

61. Hou JM, Krebs M, Ward T, et al. Circulating Tumor Cells as a Window on Metastasis Biology in Lung Cancer. American Journal of Pathology 2011; 178(3): 989–96.

62. Markidis S, Chien SWD, Laure E, Peng IB, Vetter JS. NVIDIA Tensor Core Programmability, Performance & Precision. Ieee Sym Para Distr 2018: 522–31.

63. Capra M, Peloso R, Masera G, Roch MR, Martina M. Edge Computing: A Survey On the Hardware Requirements in the Internet of Things World. Future Internet 2019; 11(4).

64. Shao C, Liao CP, Hu P, et al. Detection of live circulating tumor cells by a class of near-infrared heptamethine carbocyanine dyes in patients with localized and metastatic prostate cancer. PLoS One 2014; 9(2): e88967.

65. Duffy DC, McDonald JC, Schueller OJ, Whitesides GM. Rapid Prototyping of Microfluidic Systems in Poly(dimethylsiloxane). Anal Chem 1998; 70(23): 4974–84.

